# Signatures of mitonuclear coevolution in a warbler species complex

**DOI:** 10.1101/2020.04.06.028506

**Authors:** Silu Wang, Madelyn J. Ore, Else K. Mikkelsen, Julie Lee-Yaw, David P. L. Toews, Sievert Rohwer, Darren Irwin

## Abstract

Mitochondrial (mtDNA) and nuclear (nDNA) genes interact to govern metabolic pathways of mitochondria. When differentiated populations interbreed at secondary contact, incompatibilities between mtDNA of one population and nDNA of the other could result in low fitness of hybrids. In northwestern North America, two hybridizing species of warblers, *Setophaga occidentalis* (abbreviated as SOCC) and *S. townsendi* (STOW), provide an excellent system to investigate the potential co-adaptation of mitochondrial and nuclear DNA. The inland population of STOW (inland STOW) harbors mtDNA haplotype that is half a million years divergent from the SOCC mtDNA, and these populations also differ strongly in a few nDNA regions. Coastal populations of STOW (coastal STOW) have mixed ancestry, consistent with ancient hybridization of SOCC and inland STOW-like population. Of the few highly-differentiated nDNA regions between inland STOW and SOCC, one of these regions (on chromosome 5) is also differentiated between coastal STOW and inland STOW, and covaries with mtDNA among coastal STOW populations. Genes in this 1.2Mb region of chromosome 5 are associated with fatty acid oxidation and energy-related signaling transduction, both of which are closely associated with mitochondrial function. This chromosome 5 region is correlated with mtDNA haplotypes both within and across sampling sites, a pattern consistent with mitonuclear co-adaptation. We show that such mitonuclear coevolution might be maintained by climate-related selection, because mitonuclear ancestry is correlated with climatic conditions among sampling sites. Together, our observation suggests climatic-associated adaptation shaping mitonuclear differentiation and introgression in this species complex.

## Introduction

Mitochondrial (mtDNA) and nuclear (nDNA) genomes co-function in maintaining critical functions that influence fitness in nearly all eukaryotes ^1–5^. Populations in different areas may harbour distinct mtDNA sequences owing to selection or drift, and because many nuclear genes encode proteins that function within mitochondria, the genomes are expected to co-evolve, each being the target of selection favoring compatibility with the other ^4,6,7^. Interbreeding at species boundaries can lead to sub-optimal mitonuclear combinations in hybrids. Specifically, hybrids with nDNA from one species and mtDNA from the other species may have lowered fitness ^8,9^. These types of genetic incompatibilities can play a role in keeping hybrid zones narrow and limiting gene flow between species ^10,11^. Hence coadaptation of mtDNA and nDNA is increasingly recognized as being important to speciation ^10–12^.

In at least some species, geographic variation in climate is known to select for different mitochondrial genotypes in different areas ^6,13–15^. This may in turn lead to indirect selection on co-functioning nuclear genes. Such climatic mitonuclear coadaptation can lead to genomic differentiation between populations inhabiting different climatic conditions ^4,6^. Here we examine the relationship between mtDNA and nDNA variation in a warbler species complex with ancient and ongoing hybridization. In particular, we ask whether there is a signature of mitonuclear coevolution and, if so, could climate-related selection have driven such coevolution?

While secondary contact between differentiated populations sometimes leads to narrow hybrid zones ^16^, another possible outcome is the formation of a hybrid or mixed population over a broad region ^17–19^. Such populations have the potential to reveal strong selection on suboptimal combinations of genes from the two parental species. Despite increasing interest in mitonuclear interactions at species boundaries of natural populations with complex population histories ^3,6,20–23^, the degree to which mitonuclear interactions are important in the differentiation among lineages is not well understood.

The North America warbler species *Setophaga occidentalis* (abbreviated as SOCC) inhabits conifer forests along the states of Oregon, California, and southern Washington, U.S.A. To the north of SOCC, a closely related species *Setophaga townsendi* (abbreviated as STOW) consists of an inland population that inhabits areas east of the Coast Mountains of British Columbia, Canada and northern Washington, USA (abbreviated as inland STOW) and a coastal population (abbreviated as coastal STOW) west of the Coast Mountains (Figure 1 A) ^24–28^. The SOCC and inland STOW populations demonstrate distinct plumage and mtDNA haplotypes that are separated by ~0.8% sequence divergence, diverged ~0.5 million years ago (Figure 1 BC) ^24,26,27,29^. They differ in nDNA mainly at a few small genomic regions, one related to plumage differences (ASIP-RALY gene block), whereas the rest of the genome shows very little differentiation ^29^.

**Figure 1.**
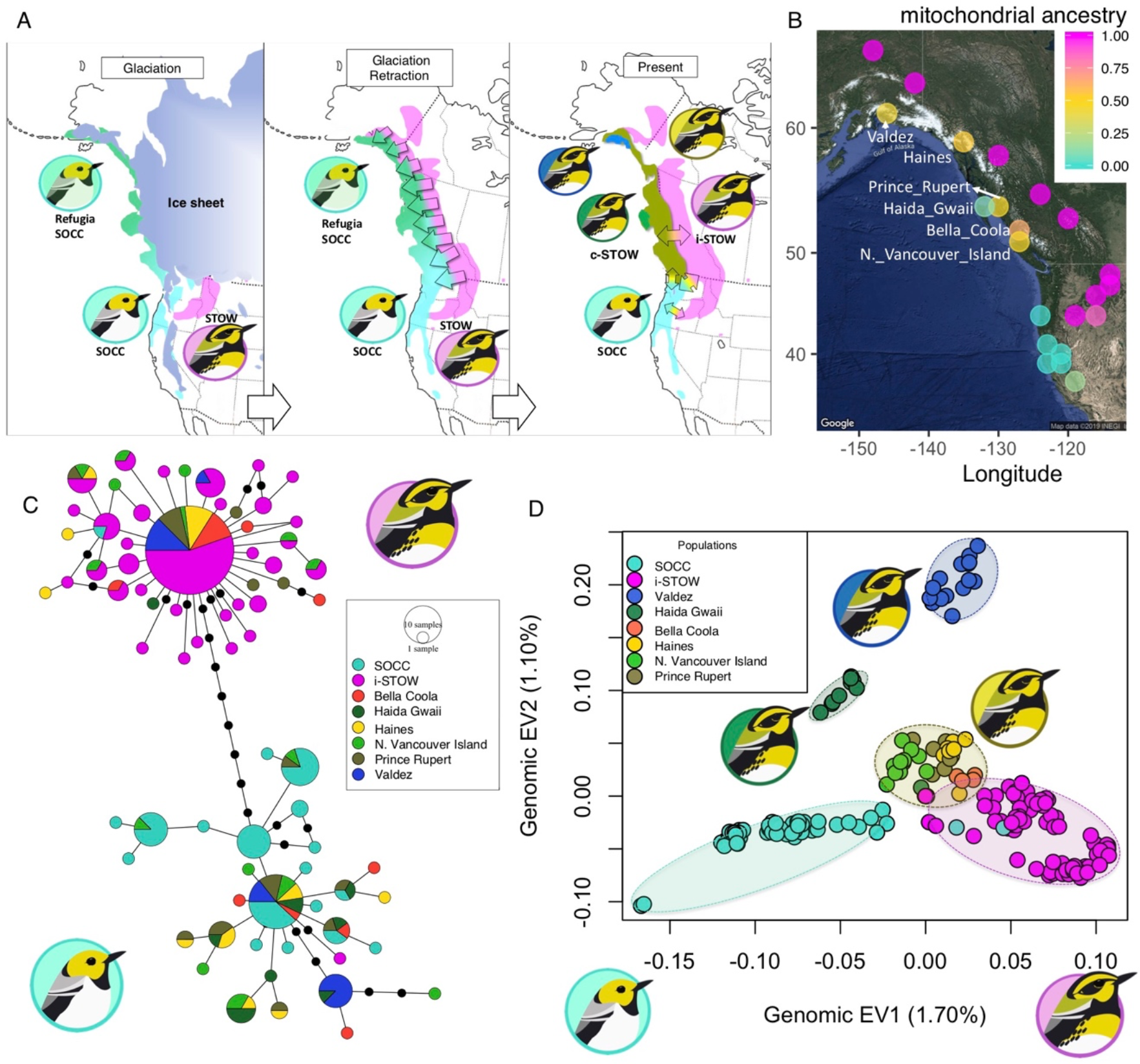
Coastal populations of *S. townsendi* (coastal STOW) demonstrate a mixture of mitochondrial and nuclear ancestry from *S. occidentalis* (SOCC) and inland *S. townsendi* (inland STOW). (**A**) Illustration of the inferred history of differentiation and hybridization between STOW and SOCC during glacial expansion and retraction. Left: During the last glacial maxima, the SOCC and STOW populations resided in isolated glacial refugia. Center: after glacial retraction, the refugial SOCC and inland STOW expanded and hybridized along a broad inland-to-coastal front parallel to the coast. Right: the historical hybridization resulted in coastal STOW populations with admixed ancestry although the plumage resembles that of inland STOW. Population substructure within coastal STOW could be a result of refugial isolation. **(B)** Mitochondrial ancestry of major sampling sites, with colour representing the proportion of individuals belonging to the two major clades shown in: (**C**) Haplotype network of mitochondrial NADH gene sequences from ref. ^27^, with colours representing major sampling areas. Each circle represents a haplotype and area of the circles are proportional to the number of individuals carrying each haplotype. The lines (regardless of their lengths) between the circles represent one mutation between haplotypes, the black dots on the lines represent additional mutations among haplotypes. SOCC and inland STOW almost universally belong to divergent mtDNA clades, whereas coastal STOW populations harbor a mixture of these two major clades. (**D**) Principle component analysis of covariance with 222,559 high quality SNPs in the nuclear genome. Populations of coastal STOW are intermediate in Eigenvector (EV) 1 but distinct from inland STOW and SOCC in EV2. EV1 represents admixture between inland STOW and SOCC. Three distinct coastal STOW clusters are evident: Valdez (royal blue), Haida Gwaii (dark green), and other coastal STOW (thereafter abbreviated as ocoastal STOW, gold).

The coastal STOW population is identical to inland STOW in plumage, but harbors both SOCC and inland STOW mtDNA haplotypes (Figure 1 BC). The haplotype network (Figure 1 C) suggests that this population is the product of ancient hybridization between SOCC and inland STOW-like population ^27^ (Figure 1 A). If so, individuals sampled from coastal STOW should demonstrate a mixture of ancestry between contemporary SOCC and inland STOW, especially detectable in regions that are differentiated between SOCC and inland STOW ^30^. In addition, the plumage gene region that differs between SOCC and inland STOW is expected to remain undifferentiated between coastal STOW and inland STOW, because of their similar appearance. In contrast, we might expect differentiation within coastal STOW with respect to nuclear genes associated with mitochondrial function according to differences in the frequency of inland STOW versus SOCC mtDNA haplotypes in different parts of the range.

Here, we analyze variation at tens of thousands of single nucleotide polymorphisms (SNPs) throughout the nuclear genome of various SOCC, coastal STOW and inland STOW populations. In particular, we ask (1) whether both nuclear and mitochondrial genomic data suggest hybrid origin of coastal STOW; (2) whether genomic differentiation between coastal and inland STOW is related to mitonuclear coevolution; (3) whether spatial variation in the frequency of mitonuclear genotypes is consistent with climate-related selection.

## Methods

### Museum samples, mtDNA sequences, and nDNA sequencing

As a baseline for understanding the relationships among mtDNA haplotypes and their distributions, sequences of the mtDNA NADH dehydrogenase subunit 2 gene (ND2) for 223 individuals (95 coastal STOW, 81 inland STOW, and 47 SOCC) from the Krosby and Rohwer (2009) ^27^ study were acquired from GenBank (accession numbers FJ373895-FJ374120). To further understand the relationships among various STOW populations with these mtDNA sequences, we generated a minimum spanning haplotype network ^31^ with PopART ^32^. This network showed two clearly separated haplotype clusters. We then scored each haplotype as 0 and 1 respectively for those that are nested within the SOCC haplotype cluster and the inland STOW cluster ^27^ (Figure 1 C).

Among these individuals with previously-sequenced mtDNA (i.e., from ^27^), we selected a subset of tissue samples (70 inland STOW, 57 coastal STOW, and 15 SOCC; obtained from the Burke Museum of Natural History and Culture, University of Washington, Seattle, Washington) for nuclear genomic sequencing. We supplemented this set of genetic samples with 54 blood samples that we obtained directly from birds caught in the field during the breeding season of 2016; these included 47 SOCC from California, 5 inland STOW from Montana, and 2 inland STOW from Washington. Therefore, 196 samples (62 SOCC, 77 inland STOW, and 57 coastal STOW) were included in the nuclear genomic sequencing.

### GBS pipeline

We prepared genotyping-by-sequencing (GBS) ^33^ libraries from DNA samples of the 196 individuals described above as our previous study ^34^. Briefly, we digested genomes with the restriction enzyme PstI, then ligated fragments with barcode and adaptors, and amplified with PCR. Amplified DNA was pooled into two libraries which were then paired-end sequenced at GenomeQuebec: the first (115 individuals) were sequenced with an Illumina HiSeq 2500 automated sequencer (read length = 125bp), and the second (81 individuals) were sequenced with an Illumina HiSeq 4000 (read length = 100bp) due to equipment updates at the sequencing facility. We randomly assigned samples to different plates and included replicates of three samples among plates. Sequence processing was based on that of a previous study ^34^, but with some changes included in the summary below. Specifically, the reads were demultiplexed with a custom script and then trimmed using Trimmomatic ^35^ [TRAILING:3 SLIDINGWINDOW:4:10 MINLEN:30]. We aligned reads to a congener *Setophaga coronata* reference ^36^ with bwa-mem ^37^ (default settings). Variable sites were identified with GATK ^38^, then filtered with VCFtools ^39^ according to the following criteria: 1) removing indels; 2) keeping sites with genotype quality (GQ) > 20; 3) keeping sites with minor allele frequency (MAF) ≥ 0.05; 4) removing sites with > 30% missing genotypes among individuals; and 5) keeping biallelic single nucleotide polymorphisms (SNPs) only. Thereafter 222,559 SNPs remained.

### Population structure and genomic differentiation

To investigate if the coastal STOW is a product of genetic admixture of SOCC and inland STOW, we first examine population structure with principle component analysis (PCA) in the SNPRelate ^40^ package in R ^41^, followed by heterozygosity-ancestry proportion analysis ^42^. We originally set out to assess the differences between SOCC, coastal STOW, and inland STOW. However, the PCA revealed strong clustering within coastal STOW with Valdez, AK (USA) and Haida Gwaii, BC (Canada) populations distinct from the rest of the coastal STOW populations. In subsequent analysis, we thus compared each of the three coastal STOW groups (Haida Gwaii, Valdez, and other coastal STOW [oc-STOW]) to the inland STOW and SOCC groups. To examine population differentiation across the genome, for each of the 222,559 filtered SNPs we calculated *F_ST_* ^43^ with VCFtools ^39^ between 1) inland STOW (*n* = 77) and SOCC (*n* = 62); 2) coastal STOW (*n* = 57: 10 Haida Gwaii, 15 Valdez, 32 others) and inland STOW; and 3) SOCC and each of the three coastal STOW clusters.

Ancient hybridization is expected to result in individuals with variable ancestry and reduced heterozygosity, in contrast to recent, early generation hybrids, which will be heterozygous across most loci. To uncover signatures of ancient hybridization, we calculated ancestry and inter-specific heterozygosity of the coastal STOW individuals. We consider SNPs with *F_ST_* > 0.6 (between inland STOW and SOCC) as being useful for informative parental ancestry. There were 50 SNPs distributed across chr 1, chr 1A, chr 4A, chr 5, chr 20, and chr Z. For each SNP, we assigned an ancestry score of 0 and 1 for homozygous SOCC and homozygous inland STOW variants respectively, and 0.5 for heterozygotes. We calculated ancestry (averaged ancestry score of the 50 SNPs) and heterozygosity of the 50 ancestry informative SNPs. To ensure that neutral sites besides selected genomic regions are considered, we repeated heterozygosity-ancestry analysis with a *F_ST_* threshold of 0.3, which involves 618 SNPs.

The 50 SNPs (from above) at *F_ST_* peaks between coastal STOW and inland STOW that are also consistent with the peaks between SOCC and inland STOW were considered candidate loci for further analyses. One possibility is that these loci are linked to genes that have a mitochondrial function and selection maintains their concordance with mtDNA ancestry.

### Association of mtDNA and nDNA

Coevolution between genomes is expected to lead to association between mtDNA and mt-associated nuclear genes within and among sites. If individuals with mismatched mt-nDNA genotypes are selected against, there should be an association between mtDNA and nDNA genotypes within each population. Such a force could be counteracted by random mating which breaks down the mt-nDNA association, thus strong selection is required to maintain adaptive mt-nDNA combinations within a single randomly mating population. Over time however, specific geographic regions may favor particular mtDNA variants and thus compatible nDNA variants, increasing mt-nDNA concordance among sampling sites.

To examine within-population association between mtDNA and nDNA ancestry, we conducted a genome-wide association study (GWAS) with GenABEL ^44^ in R to examine if there is association between mtDNA group (0 for SOCC type, or 1 for STOW type) and nuclear genotypes across the genome while controlling for population substructure. We used the *egscore* function that tests association between mtDNA ancestry and nuclear SNP ancestry while controlling for population substructure using the identity-by-state kinship matrix (calculated by the function *ibs*)^44^. We subsequently conducted genomic control assuming that randomly selected sites are not associated with mtDNA. To do so, we calculated inflation factor λ (accounting for both the effects of genetic structure and sample size) and used it to correct for the test statistic χ^2^ (and thus *p-values)* of each mt-nDNA association test. A *p-value* cutoff of 10^-6^ was employed to account for multiple hypothesis testing for sites across the genome as a balance of Bonferroni correction (0.05/222,559 SNPs ≈ 2.2 × 10^-7^) and dependency among linked SNPs. To examine gene function related to the loci associated with mtDNA haplotypes, we examined known protein-coding genes in vicinity of the candidate SNPs, using the *S. coronata* annotation ^36^. For each gene, we identified the *Taeniopygia guttata* homolog ^45^, which was then searched for Gene Ontology molecular and biological functions with UniProt ^46^.

Although within-population mitonuclear association could be largely prevented by random mating, a signature of mitonuclear coevolution might be captured in the mitonuclear ancestry associations among populations. We first calculated the mean mtDNA and nDNA ancestry of each site by averaging the locus-specific ancestry (0 for SOCC and 1 for inland STOW) among individuals. To examine between-population association between mtDNA and nDNA variants of interest, we employed a partial mantel test ^47^ with the *vegan* package in R to quantify the association between the among-sites (N =19) distance matrices of mtDNA ancestry and the nDNA ancestry, while controlling for spatial autocorrelation. In particular, the partial mantel test examined correlation between the among-sites distance matrix of mtDNA and that of the nDNA locus, conditioned on the among-sites distance matrix of coordinates.

### mtDNA sequence analysis

To add additional context to any potential functional interpretation of nuDNA – mtDNA interactions, we compared sequence divergence of protein coding genes in full mitochondrial genomes of SOCC and STOW. We used published whole-genome data from tissues sequenced by Baiz et al. (2021) ^36^ to extract high quality (>1000X coverage) mtDNA reads (*n* = 4 SOCC haplotypes, *n* = 6 STOW haplotypes; see ^36^ for details on library preparation, sequencing methodology, and sample information). Reads were aligned to the *S. coronata* mitochondrial genome, which we annotated using the MITOS^48^ web server annotation platform. To generate individual mtDNA fasta files, we used the “mpileup” command in samtools version 0.1.18 ^49^, the “call -c” command in bcftools version 1.10.2, and the “vcf2fq” option to generate fastq files. Finally, we used the program seqtk (version 1.3; https://github.com/lh3/seqtk) to translate into fasta format. We generated sequence alignments, calculated the number of fixed SNPs between the divergent haplotypes, and determined number of fixed amino acid substitutions in Geneious (11.0.3).

### Climate analysis

To investigate whether there might be selection on mt-nDNA related to climate, we tested association of site-level mt-nDNA ancestry (the averaged site ancestry score of mtDNA, chr 5 differentiation block ancestry) and climate variation. To effectively capture annual climate variation among sites, we extracted data from 26 climate variables (Table S3) from ClimateWNA ^50^ and used PCA to describe climatic variation among sites. We computed pairwise differences between sites for a) climate based on PC1, b) climate based on PC2 values, c) geographic distance, and d) mt-nuclear ancestry. We then tested for an association between climate differences among sites and differences in mt-nDNA ancestry while controlling for geographic distance using a partial mantel test in R with 10,000 permutations.

## Results

### Population structure

The mtDNA haplotype clusters are distinct between inland STOW and SOCC, with 0.8% minimum mtDNA sequence divergence between the two clades (Krosby and Rohwer 2009; Figure 1 C). Among these substitutions, there are 5 amino acid changes: 4 within NAD2 and 1 within ATP6 (Table S1). Various coastal STOW sampling sites contain a mixture of inland STOW haplotypes and SOCC haplotypes (Figure 1 C), suggesting that these coastal STOW populations are hybrid populations between inland STOW and SOCC (Krosby and Rohwer 2009). Nuclear genomic variation as assessed through variation in the 222,559 SNPs reveals isolated clusters of SOCC and inland STOW, with additional clustering within coastal STOW. The inland STOW and SOCC form two clearly differentiated clusters differing in the first eigenvector (EV1) (Figure 1 D), and most individuals from coastal STOW have a somewhat intermediate position. However, we found substructure within coastal STOW with Valdez and Haida Gwaii forming distinct genetic clusters (Figure 1 D). Valdez differs primarily along the second eigenvector (EV2), whereas Haida Gwaii differs by a combination of EV1 and EV2. Moreover, Valdez and Haida Gwaii demonstrate comparable differentiation to the parental populations as the differentiation between the parental populations (SOCC and inland STOW), as measured by genome-wide *F_ST_* (Figure S3), possibly a result of drift due to small local population size, especially on Haida Gwaii.

Analysis of ancestry and heterozygosity further supports the ancient hybrid status of coastal STOW, as coastal STOW individuals exhibit intermediate ancestry between SOCC and inland STOW but reduced heterozygosity, suggesting many generations of crossing and backcrossing (Figure 2, S2). This pattern is in contrast to what would be expected from recent, early generation hybrids, which would appear at the top of the triangle plot. The c-STOW population falls towards the inland STOW end of the ancestry spectrum, indicating that the ancient mixed population started with more inland STOW than SOCC or that there has been more introgression from inland STOW since the original admixture. Of those divergent SNPs included in the ancestry analysis, we found SOCC-biased introgression in restricted genomic regions on chr1A, chr5, and chrZ (Figure 2 B).

**Figure 2.**
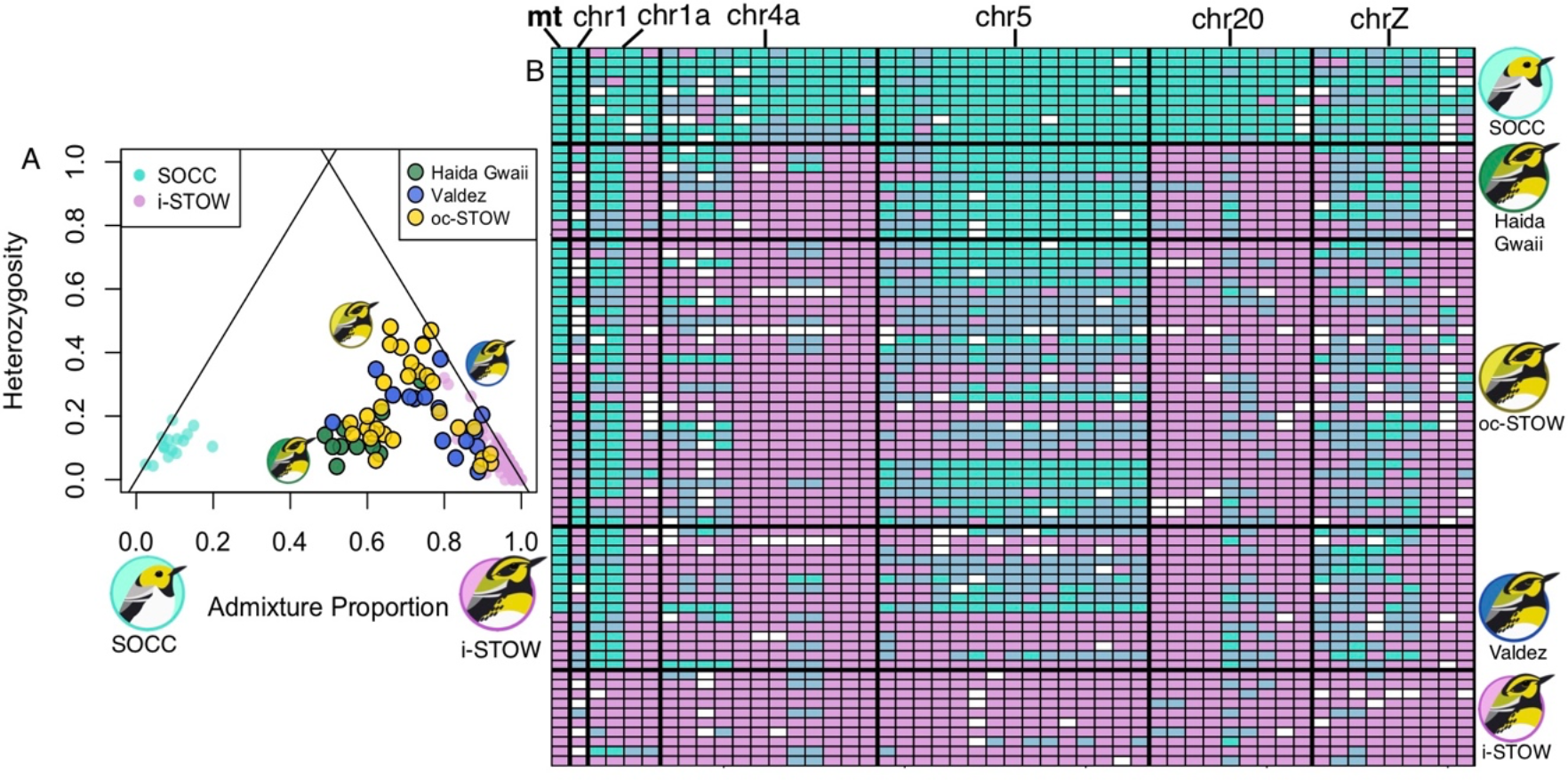
Admixture signature of coastal STOW populations. **(A)** Triangle plot showing relationship between individual admixture proportion vs. heterozygosity based on ancestry-informative SNPs (*F_ST_* > 0.6) between SOCC and inland STOW, showing that coastal STOW populations (green: Haida Gwaii, blue: Valdez; gold: oc-STOW) are ancient admixtures between SOCC and inland STOW-like population. **(B)** ancestry score (homozygotes SOCC and inland STOW respectively in turquoise and magenta, heterozygotes in light blue) of mtDNA and 50 ancestry-informative nDNA SNPs (*F_ST_* > 0.6) in chr 1, 1a, 4a, 5, 20, and Z of randomly sampled 10 individuals from parental populations (SOCC, inland STOW), and different coastal STOW populations.

### *F_ST_* distribution

Genome-wide levels of differentiation show that SOCC and inland STOW are similar (Weir and Cockerham weighted *F_ST_* = 0.023) except for a few peaks of differentiation (Figure S1 A). As in the PCA (Figure 1 D), *F_ST_* analysis indicates the Valdez and Haida Gwaii coastal STOW populations are more differentiated from both SOCC and inland STOW than other coastal STOW are (see *F_ST_* values in Figure S1, S2). The rest of the coastal STOW are more similar to inland STOW (Weir and Cockerham’s *F_ST_* = 0.008) than to SOCC (Weir and Cockerham’s *F_ST_* = 0.015) (Figure S3).

The inland STOW and SOCC have a number of peaks of differentiation (Figure S1 A) mapping to chromosomes (chr) 1, 1A, 4A, 5, 20, and Z in the *Setophaga coronata* reference. One of these (on chr 20) is in the ASIP-RALY gene block ^29^, which is known to regulate pigmentation in quail and mice ^51,52^. Our earlier study of admixture mapping in the ongoing hybrid zone between inland STOW and SOCC in the Washington Cascades ^30^ indicated that this locus is highly associated with plumage colour patterns within that zone. As predicted, the present analysis of genomic variation over a much broader geographic region shows high differentiation at the RALY SNP between sampling regions that differ in plumage (i.e., between SOCC and inland STOW, Figure S1 A, F-H) and low differentiation between regions with similar plumage (i.e., between coastal STOW and inland STOW, Figure S1 B-D).

Similar to the chr20 RALY peak, the chr5 region also showed extreme differentiation in the comparison of inland STOW and SOCC (Figure S1, 3), but in contrast to the ASIP-RALY region, the chr5 region is also strongly differentiated between coastal STOW and inland STOW and is the most differentiated region between those groups (Figure S1 B-C; 3 B-C). The chr5 peak (Figure 3 A-C) involves a ~1.2Mb region—assayed here with 15 SNPs—and includes 22 genes. These genes are predominantly associated with energy-related signaling transduction (10/22) and fatty acid metabolism (7/22) (Table S2).

**Figure 3.**
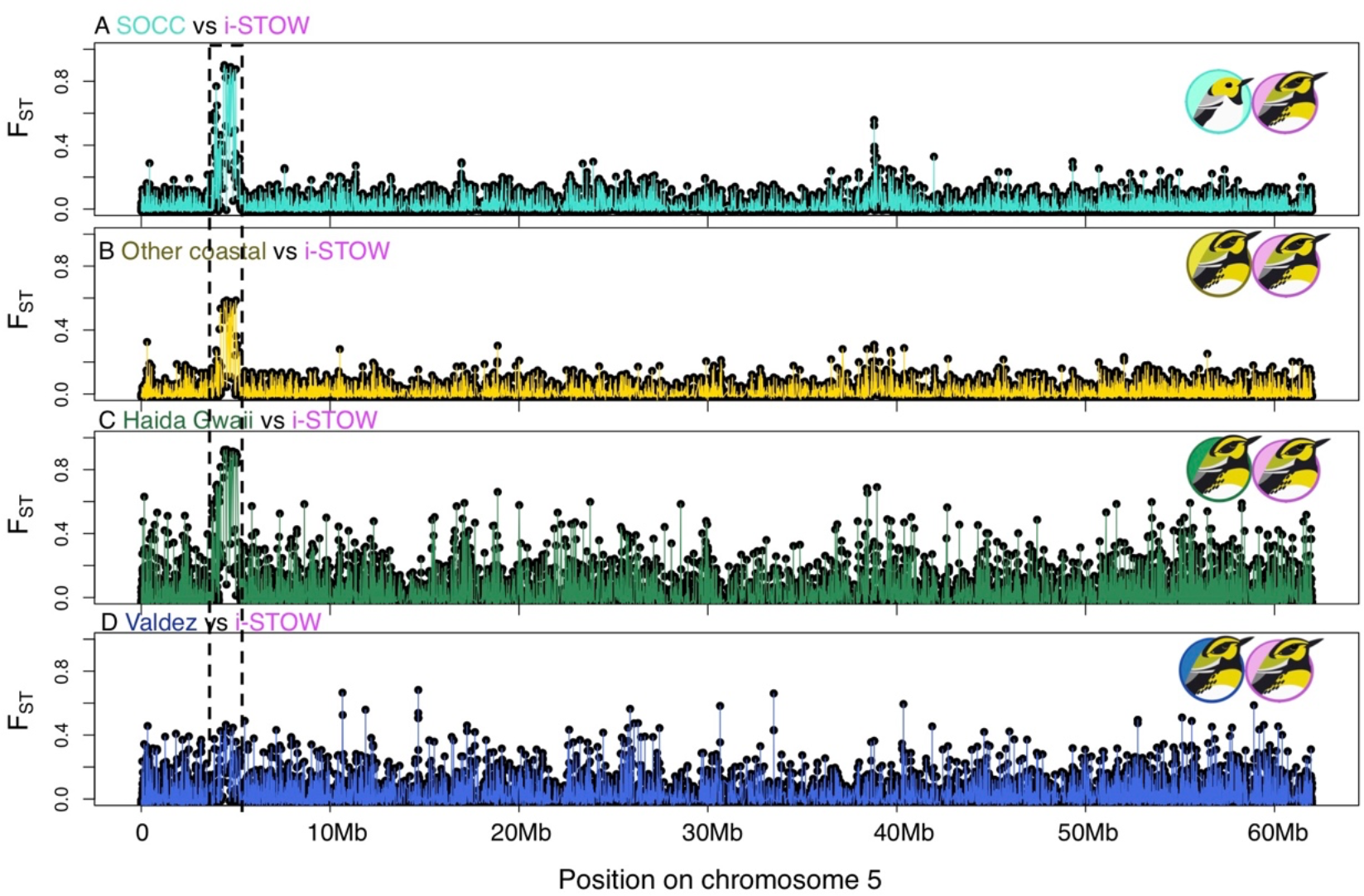
**Genetic differentiation *(F_ST_)* of variable sites along chromosome 5**, between four pairs of populations.

### Mitonuclear genetic association

We then examined whether any nuclear genomic regions were correlated with mtDNA haplotypes (Figure 1 C), controlling for population substructure. Our GWAS revealed significant *(pcorrected* < 10^-6^) mitonuclear covariation within chr 5 (Figure 4 A). The mtDNA-associated SNP on chr 5 was within the PAPLN gene (Figure 4 B, Table S2), which resides within the island of differentiation between SOCC and inland STOW (Figure 3 A; 4 B), as well as between coastal STOW and inland STOW (Figure 3, 4 B). This ~1.2Mb chr5 differentiation gene block is concentrated with genes with direct functional association with mitochondria (Table S2, energy-related signaling transduction and fatty acid oxidation). We thereafter consider this chr 5 differentiation gene block (Figure 4 C) for subsequent analysis. Among sites, the ancestry of chr 5 differentiation block (Figure 4 D, partial mantel pearson’s product-moment *r* = 0.9227,*p* < 10^-4^) were correlated with the mtDNA ancestry after controlling for spatial autocorrelation (see Methods).

**Figure 4.**
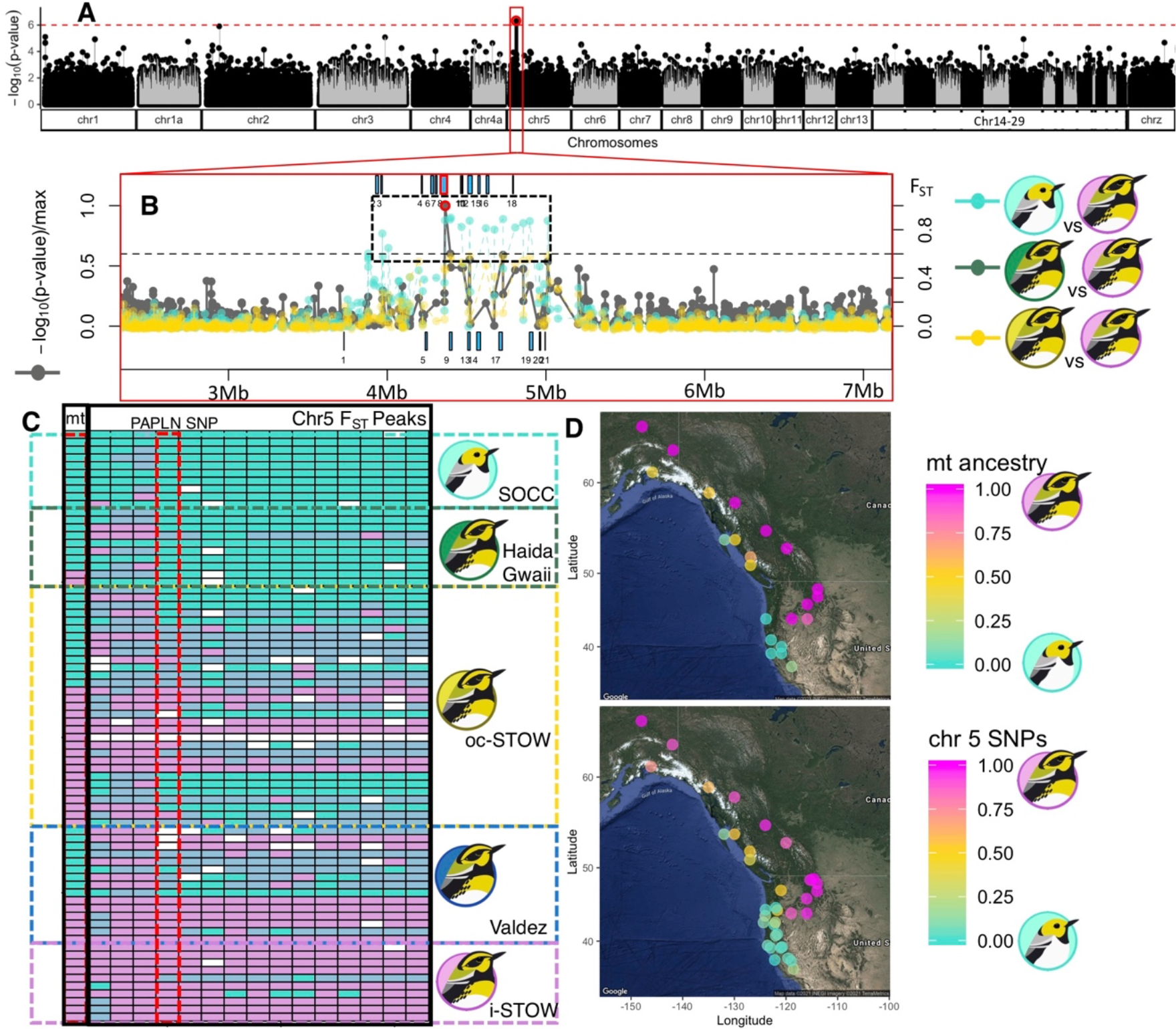
Mitonuclear association across SOCC, coastal STOW, and inland STOW populations. (**A-C**) mitonuclear association coincides with *F_ST_* peaks between SOCC and various coastal STOW versus inland STOW. (**A**) GWAS scan of mitonuclear association (controlling for population substructure) across the genome revealed significant associations *(pcorrected* < 10^-6^ indicated by red circles) within chromosome 5. (**B**) The mitonuclear association (red circle) within chromosome 5 coincides with *F_ST_* peaks (*F_ST_* > 0.6) between SOCC and inland STOW (turquoise), between Haida Gwaii STOW and inland STOW (forest green), as well as between other coastal STOW and inland STOW (yellow). Blue boxes correspond to the positions of genes (top: forward strand; bottom: reverse strand) corresponding to the numbered IDs in Table S2. The mitonuclear association peak is within Papilin, proteoglycan like sulfated glycoprotein (PAPLN) highlighted with red border. (**C**) Ancestry genotypes (turquoise = SOCC; light blue = heterozygous; magenta = inland STOW) of mtDNA and chr5 SNPs correspond to strong genotypic differences between populations. Ten randomly sampled individuals from each parental population were shown along with the individuals from various coastal STOW. The PAPLN SNP haplotypes is red bordered. (**D**) There is significant correlation between mitochondrial (top) and chr 5 (bottom) genetic block (shown in C) after controlling for spatial autocorrelation across sites (Partial Mantel Test, *p* < 10^-4^).

### Climatic association

Climate PC1 (Figure 5 AC, S5) explains 64.4% of the variation in climate among sites; this PC demonstrated similar loading among climate variables (Figure S5 AB; Table S3, S4). Climate PC2 explains 23.5% of the variation and was predominantly explained by four climate variables (Figure S5, C, S6; Table S3, S4): Temperature Difference (TD), Climate Moisture Index (CMI), Mean Annual Precipitation (MAP), Winter Precipitation (PPT_wt). The climate in coastal STOW habitat is similar to that of inland STOW along PC1, but more similar to that of SOCC along PC2, although there is great climate variation among various coastal STOW populations (Figure 5 A-C). Overall, coastal STOW habitat is moister and more stable in temperature, which is consistent with the coastal-inland humidity gradient in western North America (captured by PC2, Figure 5 B), and the distribution of mt-nDNA ancestry appears related to this geographical variation in climate. The mt-nDNA ancestry is significantly correlated with climate PC1 (Figure 5 D) (partial mantel test controlling for geographical distance, *r* = 0.307, *p* = 0.011, Figure 5 D) but not with climate PC2 (*r* = 0.166, *p* = 0.081) among 19 sites (across SOCC, coastal STOW, and inland STOW habitats).

**Figure 5.**
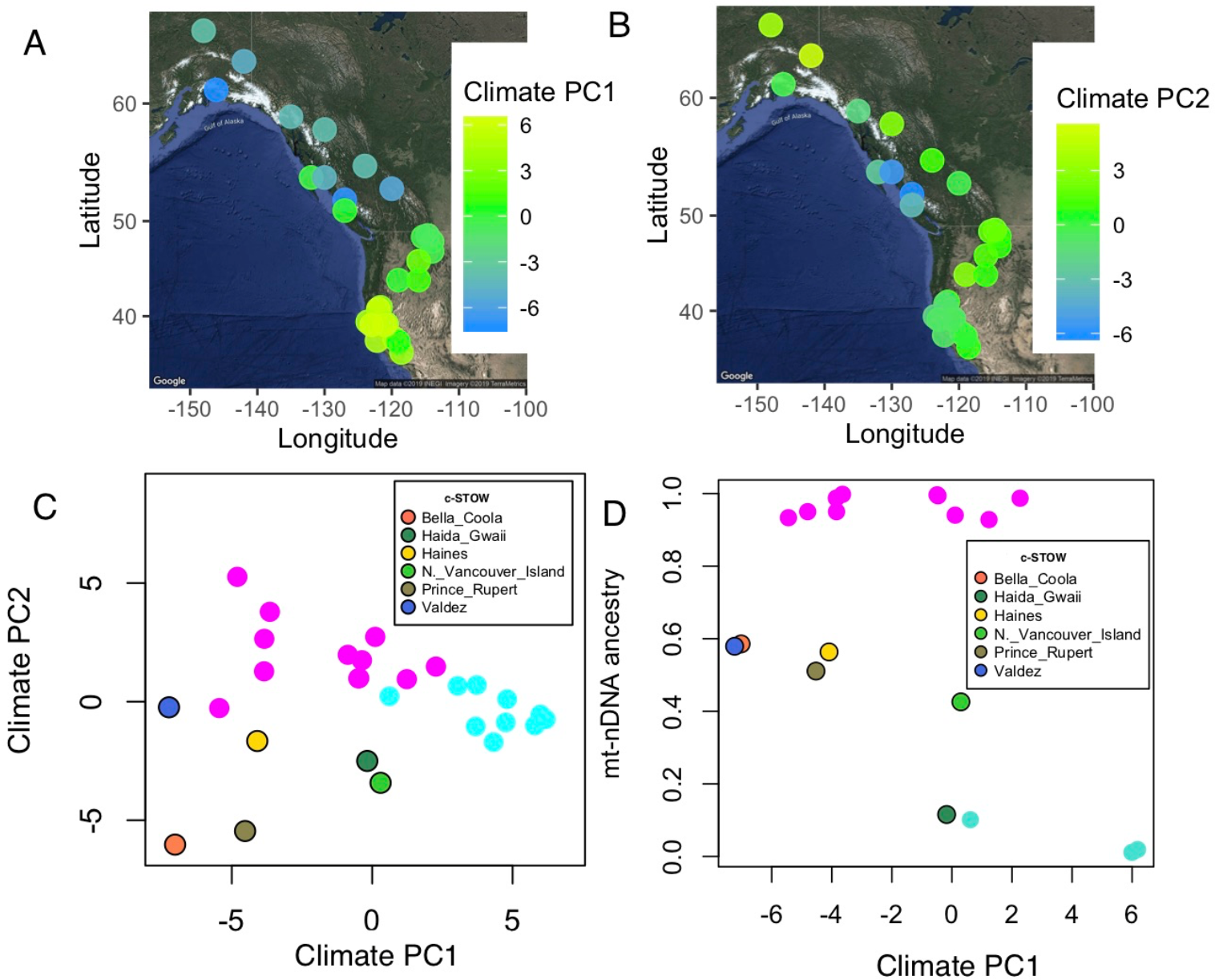
Climate principal component analysis of 26 climate variables from ClimateWNA is associated with mt-nDNA ancestry. (**A**)-(**B**), spatial variation in climate PC1 (**A**) and PC2 (**B**). (**C**) Climate PC1 and PC2, in which SOCC (turquoise), inland STOW (magenta) and coastal STOW habitats are different. (**D**) Site mean mtDNA and chr 5 gene block ancestry is correlated with local climate PC1 (partial mantel test, *r* = 0.307, *p* = 0.011).

## Discussion

*Setophaga occidentalis* (SOCC) and inland *S. townsendi* (inland STOW) are distinct in mtDNA ^27^ and exhibit three strong regions of differentiation in the nuclear genome ^30^, whereas coastal *S. townsendi* (coastal STOW) harbor admixed mtDNA and nDNA ancestry from SOCC and inland STOW. One of the regions of strong differentiation between SOCC and inland STOW, a ~1.2Mb region on chr 5, differentiates coastal STOW and inland STOW as well. This region shows strong association with divergent mtDNA haplotypes, after controlling for background population substructure of nDNA, and contains genes with functions that are involved in energy-related signaling transduction and fatty acid metabolism, both of which are strong candidates for cofunction with mtDNA. We found site-level mitonuclear ancestry covaries with the site climate conditions, consistent with adaptation to climate directly, or to other factors that are associated with climatic variation, such as forest characteristics.

### Coevolution of mtDNA and nDNA

We found the key nDNA differences between coastal STOW versus inland STOW resides at loci within chr 5 associated with fatty acid metabolism and energy-related signaling transduction (Figure 4 B; Table S2), an intriguing result given that coastal STOW and inland STOW differ so strongly in their mitochondrial haplotype frequencies. This chr5 gene block is concentrated with genes encoding proteins directly involved ATP and NADH activities. Notably, an analysis of the amino differences between whole mitochondrial genomes in this system revealed four amino acid substitutions in the *ND2* gene and one in *ATP6* (Table S1). In addition, the chr5 gene block harbors genes associated with fatty acid oxidation (Table S2), which mainly occurs in mitochondria although it occurs in other cellular compartments as well (see reviews ^53,54^). Together, these findings point to the possibility of selection on mitonuclear cofunctions ^11,12^. The SOCC nDNA may thus be partially incompatible with the inland STOW mtDNA in multiple functional roles, leading to selection against mismatched mitonuclear ancestries. Such selection maintaining mitonuclear concordance may be counteracted by random mating in admixed populations at each generation and is thus difficult to detect in samples of individuals from a single population. However, mitonuclear ancestry concordance can be more easily detected through comparison of many populations. This association (Figure 4) reveals the potential selection maintaining a functionally compatible mitonuclear fatty acid metabolism over a large temporal and spatial scale.

These mitonuclear genotypes are significantly associated with climate variation among sites, suggesting potential selection on the mt-nDNA combinations related to climate or habitat (which is also associated with climate). The climate in the coastal STOW is similar to inland STOW habitat along PC1, but similar to SOCC habitat along PC2. Correlations between any two traits that have large-scale geographic variation are expected, making it difficult to confidently infer causality from such associations alone. However, the climate and habitat differences between SOCC, inland STOW, and coastal STOW are very strong, such that these differences likely cause some selective differences. These patterns are reminiscent of the *Eopsaltria australis* (Eastern Yellow Robin) system in which distinct mt-nDNA combinations are maintained between inland and coastal habitat ^6^. Fatty acid metabolic genes have also been shown to be targets of climatic adaptation in humans, within Siberian ^55^ and Greenlandic Inuit populations ^56^. Temperature ^57^ and humidity ^58^ both influence mitochondrial fatty acid metabolism ^57,58^. Chr5-mtDNA genotypes might result in functional difference in fatty acid metabolism that is adapted to specific climate (moist and stable versus dry and variable) in the breeding habitat of these warblers.

Because SOCC has apparently inhabited coastal areas for a long period of time, the SOCC mt-nDNA gene combination may be more suited for coastal habitats compared to those of inland STOW. If the SOCC mt-nDNA genotype is favored in the coastal habitats, the frequency of SOCC mt-nDNA gene combinations would tend to increase in coastal STOW populations over time. However, ongoing gene flow between inland STOW and coastal STOW would slow down or prevent such increase. The Haida Gwaii island and Valdez population could have escaped from such a balance between selection and gene flow due to their isolation from the rest of the populations respectively by the sea and mountain ranges. Another possibility is that frequency-dependent selection is maintaining long-term mt-nDNA polymorphism in the coastal STOW. Future investigation on the spatial and temporal variation of the strength of association between mtDNA and the chr5 gene block would shed light on the evolutionary forces shaping the present and future of the coastal STOW population. Further examining such mitonuclear relationship in the three hybrid zones between SOCC and inland STOW in Washington and Oregon^59^ would shed light on the prevalence of such evolutionary forces.

### Genomic architecture of differentiation

In comparisons among these populations, the distribution of *F_ST_* across the genome is consistent with the “genic” view of differentiation ^30,60,61^, in which peaks of differentiation represent genetic targets of selection (divergent selection or selection against hybrids) that are highly distinct between populations despite the rest of the genome being homogenized by gene flow ^60–62^. Despite this ‘selection with gene flow’ scenario exhibiting a similar genomic differentiation landscape as the classic ‘divergence with gene flow’ model ^61,63^, the underlying process is different. In this system, there is a known allopatric phase when SOCC and inland STOW were separated by ice sheets (Figure 1 A) ^24,27^ Genetic differentiation that accumulated in allopatry (as opposed to gradual build up at sympatry or parapatry under ‘divergence with gene flow’) can be homogenized by hybridization at secondary contact, while the climate/habitat-related genomic target (on chr 5) of selection remains differentiated.

Between coastal STOW and SOCC, there are a number of highly differentiated loci (Figure S1 F), one of which is the ASIP-RALY gene block that was found through admixture mapping to be highly associated with plumage in the narrow Cascades hybrid zone between inland STOW and SOCC ^30^. The fact that this marker has now been shown to be strongly associated with plumage both within a local hybrid zone and throughout the whole inland STOW and SOCC species complex is strong evidence for a causal link between the RALY-ASIP genetic region and plumage differences.

### Biogeography and semi-parallel introgression

In addition to providing an empirical examination of mitonuclear coevolution, the present study also helps clarify the biogeographic history of this warbler complex. Our genomic evidence is consistent with Krosby and Rohwer’s ^27^ conclusion, based on mtDNA, that coastal British Columbia and Alaska was inhabited by geographically structured SOCC populations before inland STOW expanded from inland areas and mixed with them ^27^. The SOCC and inland STOW mtDNA haplotype groups demonstrate many differences (~0.8%), whereas both are common in coastal STOW. It is unlikely that the polymorphisms in mtDNA and nDNA in the coastal STOW were caused by incomplete lineage sorting, as opposed to hybridization (Figure 1 A). In a scenario of incomplete lineage sorting, enough time would have passed following population splitting for both inland STOW and SOCC to have lost the alternative haplotype, while the coastal STOW maintained both. Over such a period of time, sizeable differences would be expected between the SOCC haplotypes found in SOCC population versus coastal STOW, as well as between inland STOW haplotypes found in coastal versus inland STOW population. We did not observe such a pattern, as coastal STOW has some mtDNA haplotypes that are identical to SOCC and some that are identical to inland STOW haplotypes. In addition, the reduced heterozygosity (relative to expected values of recent hybrids) and intermediate ancestry (Figure 2 S2) reflected in the nuclear genome of coastal STOW further supported the ancient hybrid status of coastal STOW.

The higher genome-wide differentiation of the Haida Gwaii and Valdez populations (Figure S3) is consistent with at least partially isolated cryptic refugia of SOCC in coastal Alaska and Haida Gwaii during the last glacial maximum ^64^ Following expansion of inland STOW from the inland area, presumably after the last glacial period, hybridization between inland STOW and SOCC apparently led to populations of mixed ancestry along the coast of British Columbia and Alaska (Figure 1 A). These coastal populations have the plumage patterns and colors of inland STOW, which is why they have been classified as members of that species. This uniform inland STOW appearance has concealed a more complex history of hybridization with ancient and geographically differentiated populations of SOCC.

Following expansion of inland STOW from the interior, gene flow into Haida Gwaii may have been weak due to the expanse of water separating it from the mainland, explaining why that population is more similar to SOCC than other coastal STOW are. Gene flow into Valdez could have also been impeded by geographical barriers, as Valdez is surrounded by mountain ranges (Chugach mountains, Wrangell mountains, and St. Elias mountains). However, both nuclear and genomic data indicate that Valdez has substantial ancestry from both inland STOW and SOCC. Despite genome-wide differentiation among these three coastal STOW genetic clusters, there is an interesting parallelism: all the three populations exhibit the inland STOW-like ASIP-RALY marker (Figure S1) that is associated with plumage ^30^, and disproportionately higher frequencies of the SOCC-like chr5 nDNA gene block that covaries with mtDNA (Figure 3). Such parallelism might be driven by parallel adaptation to the coastal climate.

### Caveats and future directions

While our findings are consistent with mitonuclear coadaptation being important in the pattern of genomic differentiation within this species complex, this being an observational study we cannot definitively conclude that is the case. The strong associations between geographic variation in climate, mitochondrial haplotypes, and highly differentiated regions of the genome, along with the known roles of those divergent regions in mitochondrial-associated function and the abundant evidence for mitonuclear coadaptation in other systems ^7,11,65^, add up to strong correlative evidence for mitonuclear adaptation in this case. One possibility is that the nuclear and mitochondrial loci are independently selected by the environment, without actual coevolution between the two. Future experimental study should investigate this possibility to distinguish it from actual coevolution. If there is mitonuclear coadaptation, there should be (1) an epistatic effect of mtDNA and nDNA on fatty acid metabolic phenotypes; (2) the high frequency fatty acid metabolic phenotypes (underpinned by mitonuclear epistasis) within each site should be more fit for local climate than foreign climate.

### Conclusion

Examination of genomic differentiation in this young species group has revealed patterns consistent with climate-related coadaptation among mtDNA and nDNA involved in energy-related signaling transduction and fatty-acid metabolism. Consistent with the mtDNA pattern, the coastal STOW demonstrate both mtDNA and nuclear genomic patterns consistent with ancient admixture between inland STOW and a geographically structured ancient SOCC population. Three genetic clusters of coastal STOW are characterized by a mixed genetic ancestry between the parental populations (SOCC and inland STOW), providing natural replicates for examining the role of selection in shaping genomic differentiation. These three coastal STOW clusters exhibit differentiation from inland STOW at one of the most differentiated genomic regions (on chr 5) between inland STOW and SOCC. The geographic distributions of the mitonuclear genetic combinations related to fatty acid metabolism are associated with geographic variation in climate, suggesting mt-nDNA coevolution may have occurred in response to selection for climate adaptation. Such climate-related mitonuclear selection could be an important force driving population differentiation in this species complex.

## Data Accessibility

Sequence data is deposited in GenBank SRA (accession number: PRJNA573930; ID: 573930). Secondary analytical data tables have been deposited in dryad (https://doi.org/10.5061/dryad.44j0zpc9t).

## Acknowledgements

In support of the Bird Names for Birds movement, we omitted the existing common name of *Setophaga townsendi.* We are grateful to Sharon Birks (Burke Museum) and Chris Wood (Burke Museum) for access to the tissue samples for sequencing. We thank Geoffrey E. Hill for inspiring ideas to this study. We also thank Gil Henriques for providing digital illustrations of the warblers. For helpful discussion we thank Sally Otto, Dolph Schluter, Loren Rieseberg, Graham Coop, Dahong Chen, Mike Whitlock, Andrea Thomaz, Armando Geraldes, Meade Krosby, Hernan Morales, Xinzhu Wei, Jared Grummer, and Rohit Kolora. We are grateful for research funding provided by the Natural Sciences and Engineering Research Council of Canada (grants 311931-2012, RGPIN-2017-03919 and RGPAS-2017-507830 to DEI; and PGS D 331015731 to SW); Pennsylvania State University, the Eberly College of Science, and the Huck Institutes of the Life Sciences to DPLT; and a Werner and Hildegard Hesse Research Award in Ornithology and a UBC Four Year Doctoral Fellowship to SW. For research permits we thank Environment Canada; U. S. Geological Survey; Departments of Fish & Wildlife of Washington, Idaho, and Montana; and the UBC Animal Care Committee.

## Supplementary Figures

**Figure S1.**
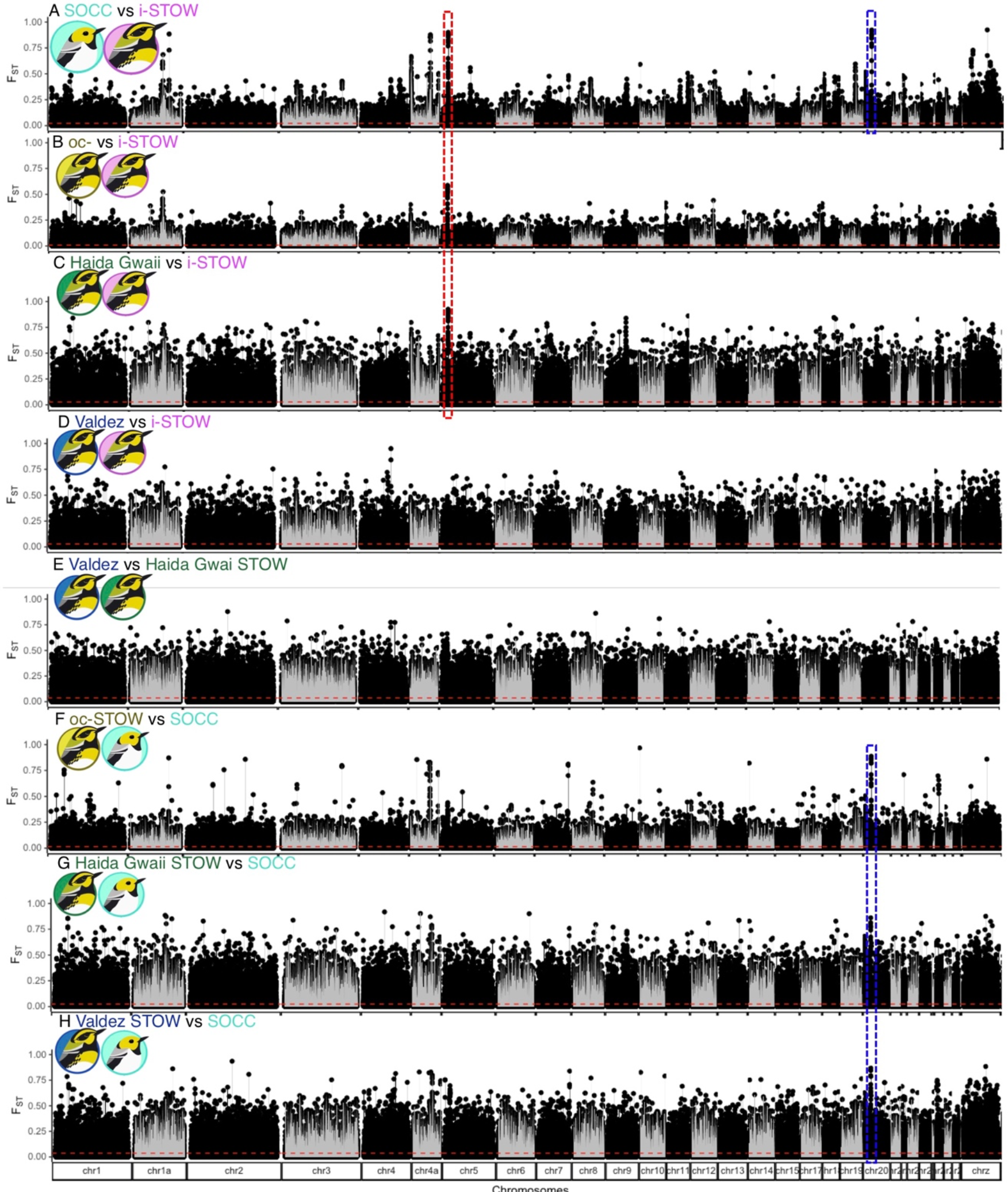
Genetic differentiation (*F_ST_*) across the genome, between pairs of *S. occidentalis* (SOCC) and/or *S. townsendi* (STOW) populations. A distinctive differentiation peak was found between coastal STOW (Haida Gwaii, Valdez, and ocoastal STOW) and inland STOW that reside in chromosome 5 (red boxes, **A**-**C**). The ASIP-RALY peak demonstrates consistent differentiation between SOCC and various STOW (blue boxes, **F**-**H**). Red horizontal dashed lines reflect Weir & Cockerham weighted average *F_ST_*.

**Figure S2.**
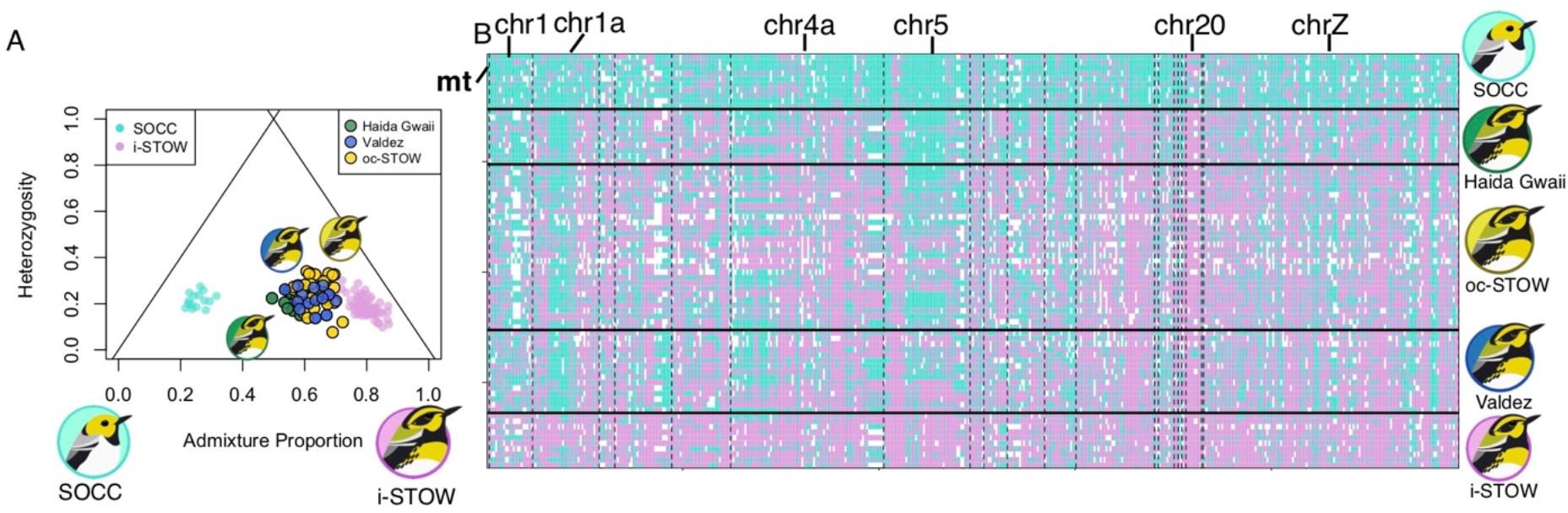
Admixture signature of coastal STOW populations. **(A)** Triangle plot showing relationship between individual admixture proportion vs. heterozygosity based on 618 SNPs (*F_ST_*>0.3) between SOCC and inland STOW, showing that coastal STOW populations consistent with ancient admixtures between SOCC and inland STOW-like population. **(B)** ancestry score (homozygotes SOCC and inland STOW respectively in turquoise and magenta, heterozygotes in light blue) of mtDNA and 618 nDNA SNPs (with *F_ST_* > 0.3) in randomly sampled 10 individuals from parental populations (SOCC, inland STOW), and different coastal STOW populations.

**Figure S3.**
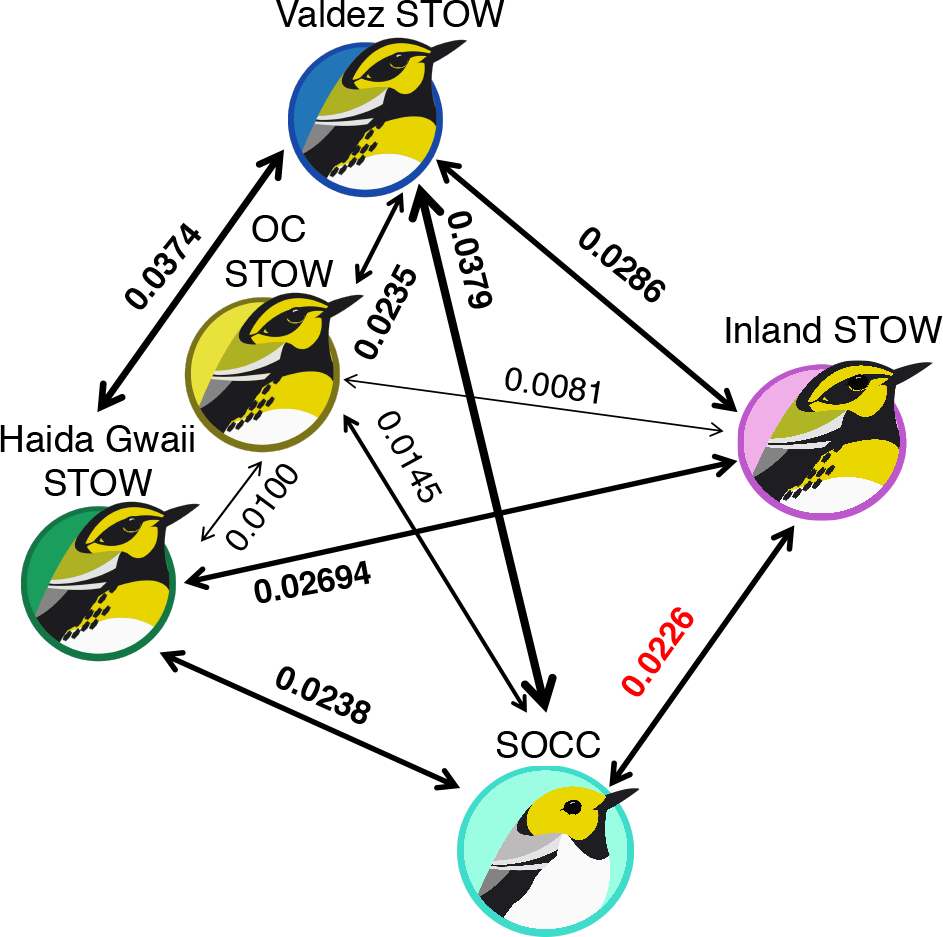
The Weir & Cockerham weighted *F_ST_* among SOCC, inland and coastal STOW. Each double-head arrow represents a pairwise comparison among the populations. The populations are oriented according to their relative geographical locations. The widths of the arrows are weighted by the *F_ST_* between each pair of populations. Surprisingly some coastal STOW populations demonstrate greater differentiation from the parental populations than between the parental populations (*F_ST_* = 0.0226).

**Figure S4.**
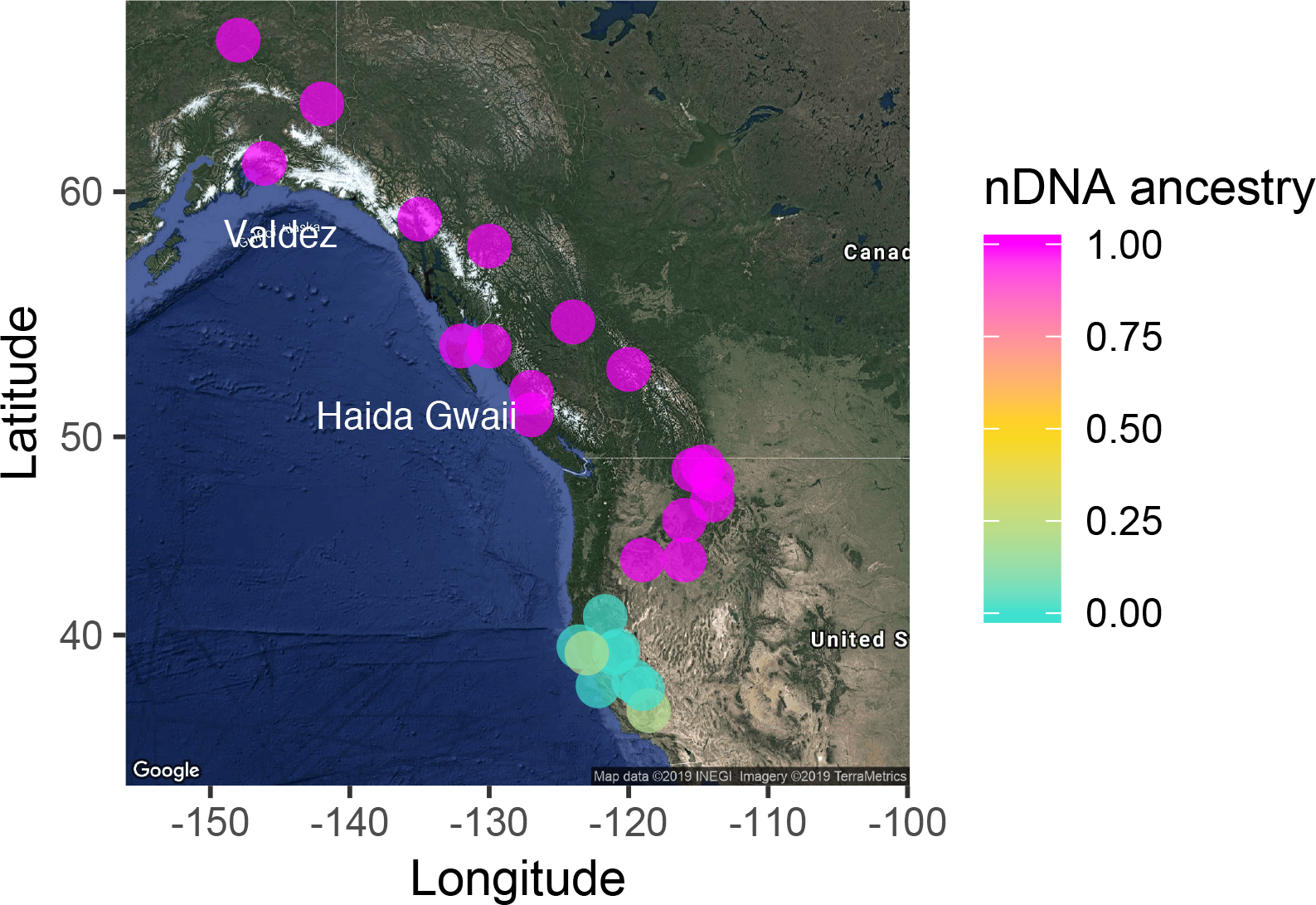
Geographical distribution of ASIP-RALY ancestry.

**Figure S5.**
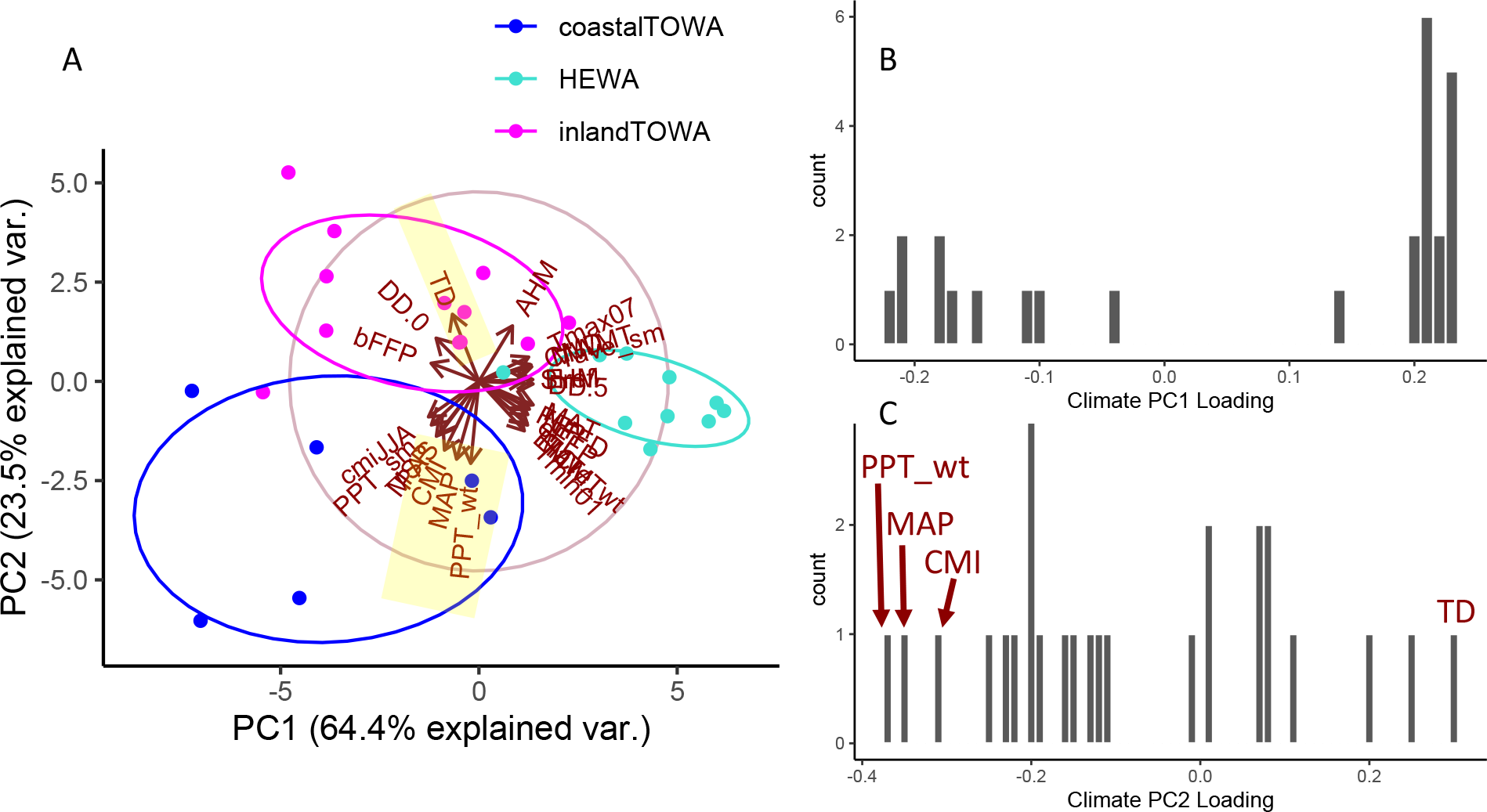
Dissecting climate PCA: **A**, biplot of PCA demonstrating loadings of the 26 climate variables in the PC space. **B**, **C**, histogram of variable loading for PC1 (**B**) and PC2 (**C**). Most of the variables demonstrates strong and even loading along PC1 (**B**), while there are 4 outstanding variables (highlighted in yellow, **A**) explaining PC2 (**C**): Temperature Difference (TD), Climate Moisture Index (CMI), Mean Annual Precipitation (MAP), Winter Precipitation (PPT_wt).

**Figure S6.**
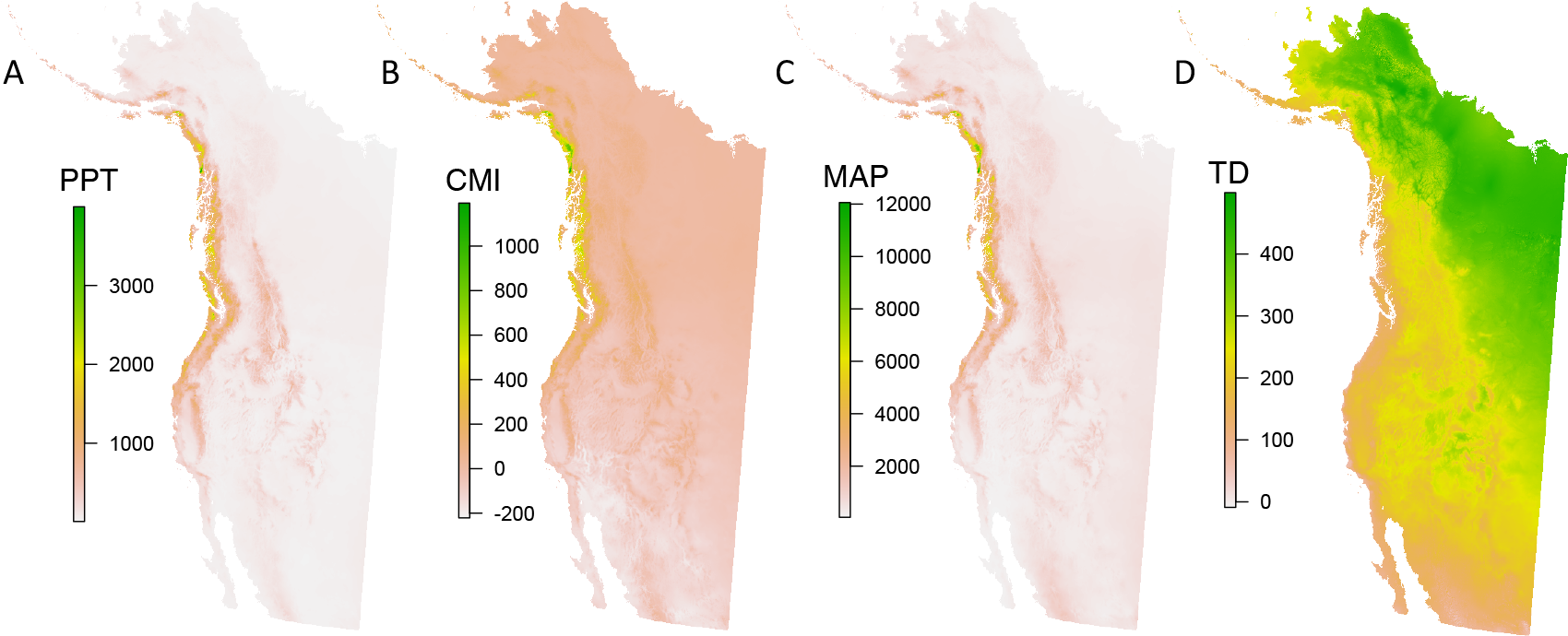
Map demonstrating spatial variation of the 4 key climate variables explaining climate PC2 (see Figure 5, S4): **A**, Winter Precipitation (PPT_wt), **B**, Climate Moisture Index (CMI), **C**, Mean Annual Precipitation (MAP), **D**, Temperature Difference (TD).

**Table S1.** Divergence of mitochondrial genes between SOCC and inland STOW.

| Name | Start | Stop | Length | %Fixed difference | Number of Amino Acid Changes |
| --- | --- | --- | --- | --- | --- |
| NAD1 | 2829 | 3776 | 948 | 0.738 | 0 |
| NAD2 | 4009 | 5043 | 1035 | 0.966 | 4 |
| COX1 | 5417 | 6949 | 1533 | 0.457 | 0 |
| COX2 | 7105 | 7779 | 675 | 0.741 | 0 |
| ATP8 | 7861 | 8022 | 162 | 0.617 | 0 |
| ATP6 | 8019 | 8699 | 681 | 0.441 | 1 |
| COX3 | 8708 | 9490 | 783 | 0.639 | 0 |
| NAD3 | 9561 | 9908 | 348 | 1.149 | 0 |
| NAD4L | 9984 | 10277 | 294 | 0.340 | 0 |
| NAD4 | 10274 | 11641 | 1368 | 0.439 | 0 |
| NAD5 | 11870 | 13660 | 1791 | 0.782 | 0 |
| COB | 13684 | 14817 | 1134 | 0.617 | 0 |
| NAD6 | 14996 | 15511 | 516 | 1.357 | 0 |

**Table S2.**
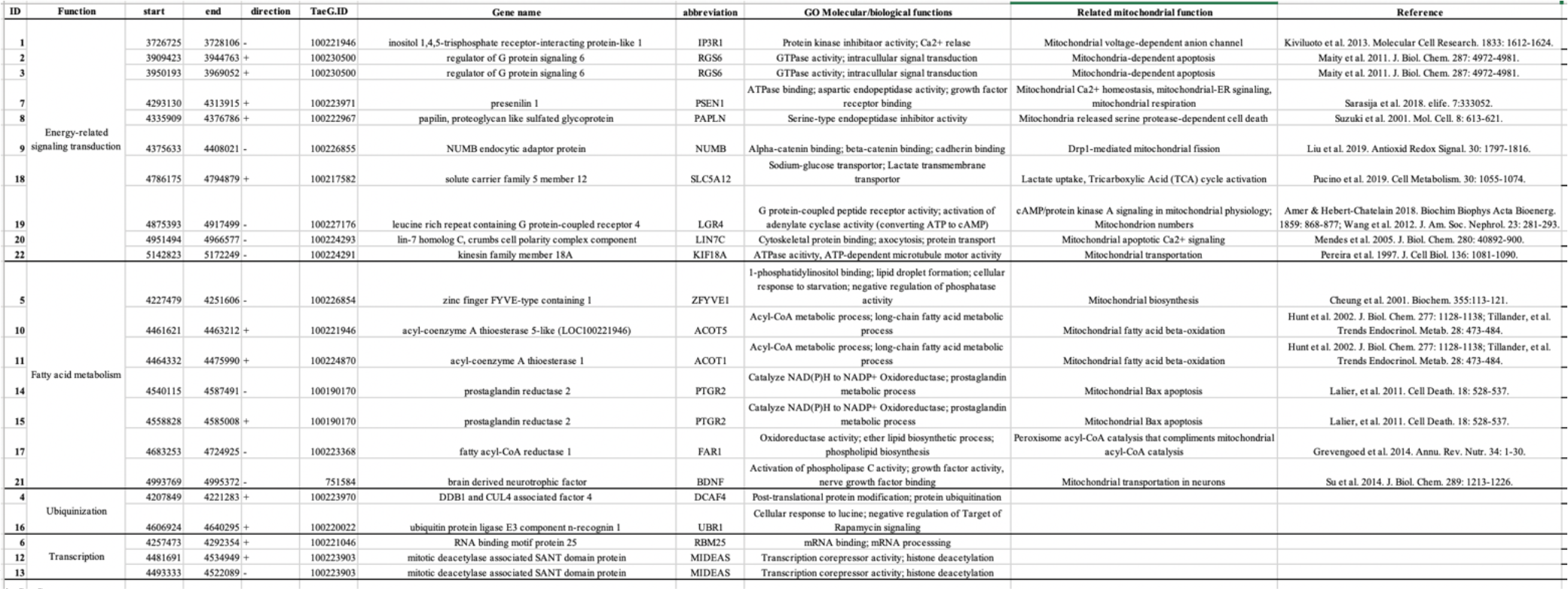
names and functions of genes involved in the 1.2 Mb chromosome 5 island of differentiation between inland STOW and coastal STOWs, as well as between inland STOW and SOCC.

**Table S3.** Definitions of climate variables from ClimateWNA ^50^.

| Column name | Variable abbreviation | Variable Name |
| --- | --- | --- |
| NORM_6190_AHM | AHM | annual heat-moisture index |
| NORM_6190_bFFP | bFFP | The day of the year on which FFP begins |
| NORM_6190_CMD | CMD | Climatic moisture deficit |
| NORM_6190_CMI | CMI | Climatic moisture index |
| NORM_6190_cmiJJA | cmiJJA | Hogg's summer (June-Aug) climate moisture index |
| NORM_6190_DD.0 | DD.0 | degree-days below 0°C, chilling degree-days |
| NORM_6190_DD.5 | DD.5 | degree-days above 5°C, growing degree-days |
| NORM_6190_eFFP | eFFP | The day of the year on which FFP begins |
| NORM_6190_EMT | EMT | Extreme minimum temperature over 30 years |
| NORM_6190_Eref | Eref | Hargreaves reference evaporation (mm) |
| NORM_6190_FFP | FFP | Frost-free period |
| NORM_6190_MAP | MAP | Mean annual precipitation (mm) |
| NORM_6190_MAT | MAT | Mean annual temperature |
| NORM_6190_MCMT | MCMT | Mean coldest month temperature (°C) |
| NORM_6190_MSP | MSP | May through September precipitation |
| NORM_6190_MWMT | MWMT | Mean winter temperature |
| NORM_6190_NFFD | NFFD | number of frost-free days |
| NORM_6190_PAS | PAS | precipitation as snow |
| NORM_6190_PPT_sm | PPT_sm | summer precipitation (mm) |
| NORM_6190_PPT_wt | PPT_wt | winter precipitation (mm) |
| NORM_6190_SHM | SHM | summer heat-moisture index<br>((MWMT)/(MSP/1000)) |
| NORM_6190_Tave_sm | Tave_sm | summer mean temperature (°C) |
| NORM_6190_Tave_wt | Tave_wt | winter mean temperature (°C) |
| NORM_6190_TD | TD | Continentality |
| NORM_6190_Tmax07 | Tmax07 | winter mean maximum temperature (°C) |
| NORM_6190_Tmin01 | Tmin01 | winter mean minimum temperature (°C) |

**Table S4.** The loading of variables in the climate PCA sorted by their loading along PC1.

| **Climate**<br>**Variable** | **PC1 loading** | **PC2 loading** |
| --- | --- | --- |
| cmiJJA | -0.21763162 | -0.161032306 |
| bFFP | -0.20625054 | 0.082056348 |
| PPT_sm | -0.20510277 | -0.197710506 |
| DD.0 | -0.18472653 | 0.195940506 |
| MSP | -0.183508 | -0.248663084 |
| PAS | -0.1660562 | -0.234659637 |
| CMI | -0.1486023 | -0.311869513 |
| TD | -0.11435537 | 0.30417286 |
| MAP | -0.09633503 | -0.352869482 |
| PPT_wt | -0.03530235 | -0.371886563 |
| AHM | 0.14480557 | 0.251595965 |
| Tmin01 | 0.19581554 | -0.22437128 |
| SHM | 0.19978674 | 0.007101683 |
| MCMT | 0.20705781 | -0.195744433 |
| Tave_wt | 0.20859601 | -0.192174784 |
| EMT | 0.20990053 | -0.198261928 |
| eFFP | 0.2107584 | -0.150721133 |
| FFP | 0.21127471 | -0.117676014 |
| Tmax07 | 0.21473286 | 0.110168321 |
| NFFD | 0.21782119 | -0.126257635 |
| CMD | 0.22005085 | 0.076818851 |
| MWMT | 0.22507318 | 0.074770918 |
| Tave_sm | 0.22805563 | 0.066315194 |
| Eref | 0.23258512 | 0.008062124 |
| DD.5 | 0.23376903 | -0.011649806 |
| MAT | 0.23420797 | -0.106646542 |

